# TRPV1 feed-forward sensitisation depends on COX2 upregulation in primary sensory neurons

**DOI:** 10.1101/2020.09.05.267708

**Authors:** Tianci Li, Gaoge Wang, Vivian Chin Chin Hui, Daniel Saad, Joao de Sousa Valente, Paolo La Montanara, Istvan Nagy

## Abstract

Increased activity and excitability (sensitisation) of a series of molecules including the transient receptor potential ion channel, vanilloid subfamily, member 1 (TRPV1) in pain-sensing (nociceptive) primary sensory neurons are pivotal for developing pathological pain experiences in tissue injuries. TRPV1 sensitisation is induced and maintained by two major mechanisms; post-translational and transcriptional changes in TRPV1 induced by inflammatory mediators produced and accumulated in injured tissues, and TRPV1 activation-induced feed-forward signalling. The latter mechanism includes synthesis of TRPV1 agonists within minutes, and upregulation of various receptors functionally linked to TRPV1 within a few hours, in nociceptive primary sensory neurons. Here, we report that a novel mechanism, which contributes to TRPV1 activation-induced TRPV1-sensitisation within ~30 minutes in at least ~30% of TRPV1-expressing cultured murine primary sensory neurons, is mediated through upregulation in cyclooxygenase 2 (COX2) expression and increased synthesis of a series of COX2 products. These findings highlight the importance of feed-forward signalling in sensitisation, and the value of inhibiting COX2 activity to control pain, in nociceptive primary sensory neurons in tissue injuries.

## INTRODUCTION

Tissue injuries are followed by an inflammatory reaction, which aims to restore tissue integrity (Placek *et al*, 2019). This inflammatory process is associated with pathological pain experiences including hypersensitivity to mechanical and heat stimuli known respectively as mechanical allodynia and heat hyperalgesia in human (Berta *et al*, 2017; Nagy, 2004; Yekkirala *et al*, 2017). The non-selective cationic channel, transient receptor potential ion channel, vanilloid subfamily, member 1 (TRPV1) is expressed in a major group of pain-sensing (nociceptive) primary sensory neurons, the first neurons detecting tissue pathology, and it plays a pivotal role in the development of heat hyperalgesia and significantly contributes to the development of mechanical allodynia in inflammation following tissue injuries (Caterina *et al*, 2000; Caterina *et al*, 1997).

The development of hypersensitivities depends on plastic changes, known as sensitisation (a use-dependent increase in the activity and excitability), of neurons involved in nociceptive processing (Woolf & Ma, 2007). Upregulation of TRPV1 expression, TRPV1 translocation from the cytoplasm to the cytoplasmic membrane and post-translational modification-mediated increases in TRPV1 responsiveness and activity significantly contribute to the sensitised state of nociceptive primary sensory neurons and underlay the pivotal role of TRPV1 in the development of hypersensitivities to heat and mechanical stimuli in tissue inflammation (Amadesi *et al*, 2006; Fukuoka *et al*, 2002; Ji *et al*, 2002; Kao *et al*, 2012; Khan *et al*, 2008; Moriyama *et al*, 2005; Premkumar & Ahern, 2000; Rathee *et al*, 2002; Van Buren *et al*, 2005; Zhang *et al*, 2005). Inflammatory mediators including nerve growth factor, bradykinin, prostaglandins (PGs), or ligands of various protease activated receptors (PARs), such as thrombin, mast cell-derived tryptase or kallikrein acting on their cognate receptors on TRPV1-expressing primary sensory neurons, initiate those changes in TRPV1 (Isensee *et al*, 2014; Ji *et al.*, 2002; Khan *et al.*, 2008; Moriyama *et al.*, 2005; Taiwo *et al*, 1989; Taiwo & Levine, 1990). In addition, TRPV1 activation itself may also increase its own responsiveness and activity through feed-forward mechanisms occurring at various levels of signalling in primary sensory neurons. First, TRPV1 activation through Ca^2+^ influx induces the synthesis of the TRPV1 endogenous ligand anandamide in TRPV1-expressing primary sensory neurons within minutes (Nagy *et al*, 2009; van der Stelt *et al*, 2005). Second, TRPV1 activation also induces, within hours, upregulation of the expression of genes whose products are implicated in post-translational modification-mediated increases in the responsiveness and activity of TRPV1 (Chen *et al*, 2013). Recently, Zhang and colleagues have reported a third type of feed-forward TRPV1 sensitisation that develops within ~ 20-30 minutes following TRPV1 activation by the archetypical exogenous ligand capsaicin and is mediated through extracellular signal-regulated kinase 1/2 (ERK1/2) and the Ca^2+-^ calmodulin-dependent protein kinase II (CaMKII) activity (Zhang *et al*, 2011). Here, we show that this third type of feed-forward TRPV1 sensitisation depends on upregulation of cyclooxygenase 2 (COX2) and the subsequent increase in the synthesis of a series of PGs.

## RESULTS

### Capsaicin induces sensitisation of responses to subsequent capsaicin application in ~30 minutes in mouse cultured primary sensory neurons

The most consistent development of sensitised responses to second capsaicin application was observed when 500nM capsaicin, which is about the EC50 of this compound at mouse TRPV1 (Caterina *et al.*, 2000), was applied for 15 seconds 30 minutes before the second capsaicin (500nM) application. The 30 minutes superfusion of the cells with the extracellular buffer however, resulted in a significant reduction in KCl-evoked responses indicating a general “run down” of the responses (p<0.0001, Student’s t-test, n=288; Figure 1A-C; Supplementary Figure 1). Therefore, the amplitude of each capsaicin-evoked response was normalised to the amplitude of the subsequent KCl-evoked response. Then, the first and second normalised capsaicin-evoked responses were compared. This comparison revealed that, in 84 of 288 neurons, the second normalised capsaicin-evoked response was more than 5% greater than the first response (Figure 1A and B). These cells were considered sensitizer neurons. All other neurons (n=204) were considered as “non-sensitizer” (Figure 1C). Within the 84 sensitizers, three cells were considered outliers as the ratio of the first and second capsaicin-evoked responses significantly exceeded (11.84, 20.63 and 120.41) the average of the 84 cells (3.035±2.44; n=84). These three outliers were not included in the statistical analysis of the control recordings.

**Figure 1.**
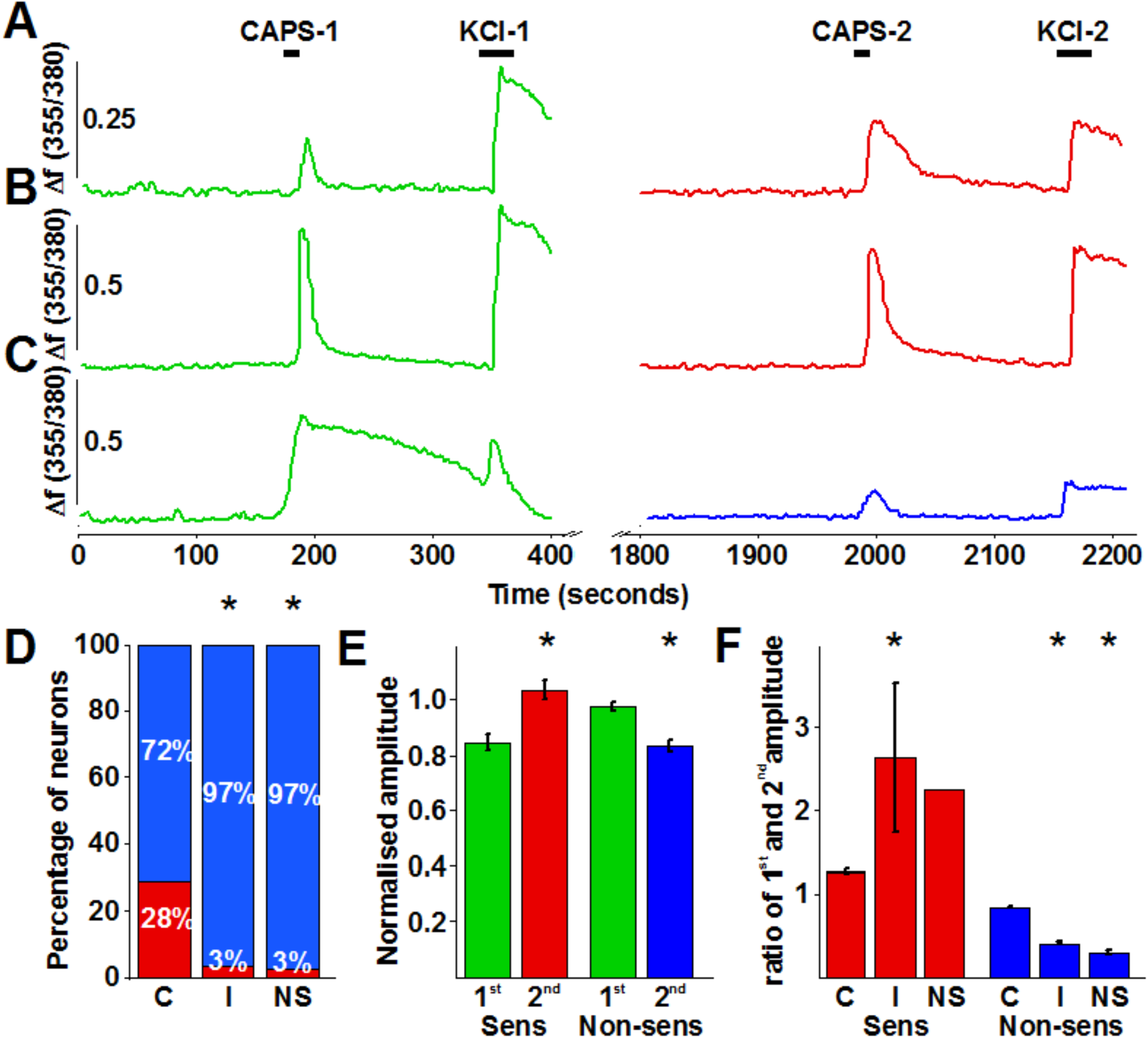
Capsaicin application to cultured murine primary sensory neurons induces sensitised responses to capsaicin applied 30 minutes after the first application in a group of neurons in a COX2-dependent manner. (A), (B) and (C) shows typical recordings of changes in the intracellular Ca2+ concentrations in three cultured murine primary sensory neurons in control extracellular buffer. Capsaicin (500nM) was applied for 15 seconds at 3 (CAPS-1) and 33 minutes (CAPS-2). Applications of capsaicin were followed by the applications of KCl (50mM) 2 minutes after capsaicin applications (KCl-1 and KCl2). The neurons whose responses are shown in (A) and (B) exhibited absolute (A), or relative to the KCl-evoked response (B), sensitised second capsaicin-evoked responses when the amplitudes are assessed. The cell whose response is shown in (C) exhibited a non-sensitised response. (D) Relative number of neurons exhibiting sensitizer (red) or non-sensitizer (blue) phenotype in control extracellular buffer (C), and in the presence of indomethacin (10μM; I) or NS-398 (3μM; NS). Both indomethacin and NS-398 significantly reduced the proportion of sensitizers (p<0.0001; Fischer’s exact test for indomethacin and p<0.0015; Fischer’s exact test for NS-398). (E) Averaged normalised amplitudes (to the amplitudes of KCl-evoked responses) of first (green) and second (red for sensitizers (Sens), blue for non-sensitizers (Nonsens)) capsaicin application-evoked responses. In sensitizers and non-sensitizers the second capsaicin-evoked responses were, respectively, significantly greater (p<0.0001; Student’s paired t-test; n=81) and smaller (p<0.0001, Student’s paired t-test; n=204) then the amplitudes of the first capsaicin-evoked responses. (F) Ratio of first and second normalised (to KCl-evoked responses) capsaicin-evoked responses in control buffer (C) and in the presence of indomethacin (10μM; I) or NS398 (3μM; NS) in sensitizer (Sens) and non-sensitizer (Non-sens) neurons. Both indomethacin and NS-398 increased or reduced the ratio in sensitizer or non-sensitizer neurons, respectively (indomethacin: p<0.0001; Student’s t-test for sensitizers and p<0.001; ANOVA followed by Tukey’s post-hoc test for non-sensitizers; NS398: p<0.0001; ANOVA followed by Tukey’s post-hoc test for non-sensitizers).

On average, the normalised amplitudes of the first and second responses of the 81 sensitizers were 0.85±0.03 and 1.04±0.04 (p<0.0001; Student’s paired t-test; n=81), whereas those for the non-sensitizer cells were 0.98±0.02 and 0.83±0.02 (p<0.0001; Student’s paired t-test; n=204; Figure 1E). Both the first and second average responses in the sensitizer (n=81) and non-sensitizer neurons (n=204) were significantly different (first responses: p=0.0002, Student’s two samples t-test; second responses: p<0.0001, Student’s two samples t-test). The normalised first and second responses of both the sensitizer and non-sensitizer cells exhibited correlation (R=0.85, p<0.0001 for the sensitizers and R=0.77, p<0.0001 for the non-sensitizers; Supplementary Figure 2A).

The average ratio of the first and second capsaicin-evoked responses was 1.26±0.03 (n=81) in the sensitizer, while it was 0.84±0.01 (n=204) in the non-sensitizer group (p<0.0001, Student’s two samples t-test; Figure 1F). Within the sensitizers, cells with smaller normalised first responses tended to show greater increase (R=-0.49, p<0.0001; Supplementary Figure 2B), whereas non-sensitizers exhibited a very weak correlation between the amplitude of the first normalised responses and the ratio of the first and second responses (R=0.21, p=0.003; Supplementary Figure 2B).

### The development of capsaicin-induced sensitisation of subsequent capsaicin-evoked responses depends on COX2

The 30-minute delay in the development of sensitised capsaicin-evoked responses following capsaicin application suggests that the sensitisation depends on changes in the transcription of immediate early genes (Zhang *et al.*, 2011; current data). A search for immediate early gene products that could induce TRPV1 sensitisation through ERK1/2 and CaMKII, the signalling molecules shown being involved (Zhang et al., 2011), suggested that upregulation in *ptgs2* and subsequent upregulation in COX2 expression and increased synthesis of COX2 products might underlie the development of capsaicin-induced sensitisation of subsequent capsaicin-evoked responses. Therefore, next we applied indomethacin (10μM), a non-selective COX inhibitor, to the cells during the interim period between the two capsaicin applications. Indomethacin significantly reduced the proportion of cells exhibiting sensitised responses to ~3% (8 of 244; p<0.0001; Fischer’s exact test; Figure 1D). Interestingly, while the degree of sensitisation was significantly greater in the remaining sensitizers (2.63±0.9, n=8 in indomethacin and 1.26±0.03, n=81 in control; p<0.0001; ANOVA followed by Tukey’s post-hoc test; Figure 1F), the ratio of the first and second normalised capsaicin-evoked responses in the non-sensitizer cells was significantly smaller in the presence of indomethacin (0.41±0.2, n=236 in indomethacin and 0.84±0.01, n=204 in control; p<0.001; ANOVA followed by Tukey’s post-hoc test; Figure 1F).

Next, we applied the selective COX2 inhibitor NS-398 (3μM) to the cells to specifically explore whether COX2, whose products have been reported to contribute to TRPV1 sensitisation (Moriyama *et al.*, 2005; Taiwo *et al.*, 1989; Taiwo & Levine, 1990), is responsible for the sensitisation of the second capsaicin-evoked responses. As seen with indomethacin, NS-398 also significantly reduced the proportion of the sensitising neurons (~3%, 1 of 37; p<0.0015; Fischer’s exact test; Figure 1D). Again, the size of sensitisation in the single sensitizer neuron appeared greater (2.24±0.9, n=1 in NS-398), whereas the ratio of the first and second normalised capsaicin-evoked responses in the non-sensitizers was significantly smaller (0.3±0.03, n=36; p<0.0001; ANOVA followed by Tukey’s post-hoc test; Figure 1F) in the presence of NS-398. Collectively, these data suggested that capsaicin induced upregulation of *ptgs2*, COX2 and COX2 products could indeed significantly contribute to capsaicin-evoked sensitisation of subsequent capsaicin-induced responses in primary sensory neurons. Thus, we further tested this hypothesis.

### Capsaicin application to mouse cultured primary sensory neurons induces *ptgs2* and COX2 upegulation

Conventional RT-PCR was used to assess *ptgs2* expression in cultured primary sensory neurons after treating them with capsaicin or buffer for control. The size of the RT-PCR product (214 bp) was indistinguishable from the predicted size (Figure 2A; Supplementary Figure 3). Analysis of gel images revealed that capsaicin treatment indeed significantly upregulated *ptgs2* expression (p=0.0004; Student’s two samples t-test, n=3) in cultured primary sensory neurons (Figure 2A and B).

**Figure 2.**
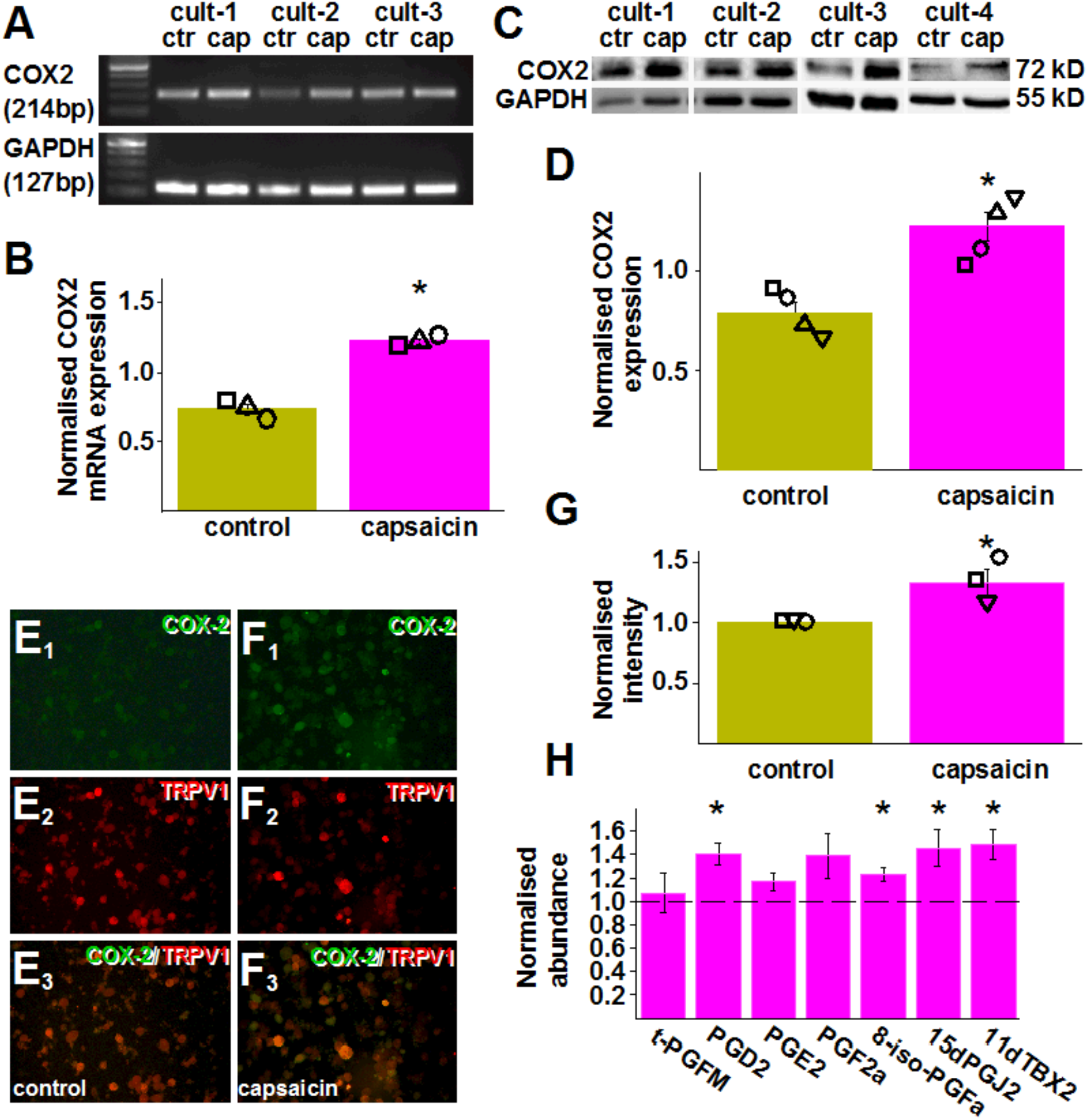
Capsaicin application to cultured murine primary sensory neurons induces upregulates COX2 expression and synthesis of a series of COX products. (A) Gel image of RT-PCR products amplified using primer pairs for murine *ptgs2* (COX2) and *gapdh* (GAPDH) and cDNA prepared using RNA isolated from three cultures (cult-1 – cult-3) of murine primary sensory neurons 25 minutes after incubating the cells in control (ctr) buffer or in the presence of 500nM capsaicin (cap) for 5 minutes. (B) Averaged normalised *ptgs2* expression in cultures shown in (A). Capsaicin (500nM, capsaicin) application resulted in a significant increase in *ptgs2* expression (p=0.0004; Student’s t-test, n=3). (C) Gel images of immunoblots prepared using proteins extracted from four cultures (cult-1 – cult-4) of murine primary sensory neurons 25 minutes after incubating the cells in control (ctr) buffer or in the presence of 500nM capsaicin (cap) for 5 minutes and antibodies raised against COX2 and GAPDH. (D) Averaged normalised COX2 expression in cultures shown in (C). Capsaicin (500nM, capsaicin) application resulted in a significant increase in COX2 expression (p=0.0025; Student’s t-test, n=4). (E_1_), (E_2_) (E_3_) (F_1_) (F_2_) (F_3_) Microscopic images of cultured murine primary sensory neurons incubated in control buffer ((E_1_), (E_2_) (E_3_)) or in the presence of 500nM capsaicin ((F_1_) (F_2_) (F_3_) for 5 minutes. Cultures were fixed 25 minutes after the incubation and incubated in anti-COX2 (green) and anti TRPV1 antibodies (red). (E_1_), (F_1_) show COX2 staining, (E_2_), (F_2_) show TRPV1 staining whereas (E_3_), (F_3_) show composite image. Note that COX2 staining intensity is increased in the culture incubated in the presence of capsaicin. (G) Normalised averaged intensity values of COX2 immunostaining in three cultures of murine primary sensory neurons. Capsaicin significantly increased the staining intensity (p=0.0337; Student t-test; n=3). (H) Targeted UPLC-MS measurements of seven COX products in an eicosanoids panel using samples prepared from cultured murine primary sensory neurons incubated in control buffer or 500nM capsaicin for 5 minutes. Samples were prepared 25 minutes after finishing the 5 minutes incubation. During the 25 minutes, cells were kept in the presence of the COX2 substrate arachidonic acid (10μM). Four of the seven COX products present in the eicosanoid panel exhibited significant increase by incubating the cells in capsaicin (PGD2: q=0.032; 8-iso-PGF2α: q=0.028; 15dPGJ2: q=0.046; 11dTXB2=0.031; Student-t test followed by Benjamini-Hochberg FDR).

Next, we used Western blotting and immunostaining to confirm that the *de novo* synthesized *ptgs2* is translated into protein. Western blot images confirmed that cultured primary sensory neurons exhibit basal COX2 expression (Figure 2C; Supplementary Figure 4; Li et al., 2011, Fehrenbacher et al., 2005, Samad et al., 2001, Araldi et al., 2013). Image analysis also revealed that capsaicin exposure induced a significant increase in that expression (p=0.0025; Student’s t-test, n=4; Figure 2C and D).

Immunostaining of control cultures also confirmed basal COX2 expression in a population of neurons (Figure 2E-F; Li et al., 2011, Fehrenbacher et al., 2005, Samad et al., 2001, Araldi et al., 2013). Capsaicin exposure significantly increased the intensity of COX2 immunopositivity (p=0.0337; Student t-test; Figure 2E–G). Importantly, about half the TRPV1-expressing cells exhibited COX2 immunopositivity in both conditions (Figure 2E-F).

### Capsaicin application to mouse cultured primary sensory neurons increases the concentration of a series of COX2 products

To confirm that the upregulated COX2 expression is associated with increased PG synthesis, we collected cells and the superfusate of mouse cultured primary sensory neurons following their exposure to capsaicin or buffer for control, and used ultra-performance liquid chromatography with tandem mass spectrometry (UPLC-MS) for targeted analysis of the samples for a group of arachidonic acid-derived compounds (Supplementary Table 1). Our measurements showed that only a few fatty acids, LOX products and one COX product were above the detection level in the first experiment (Supplementary Table 1). Therefore, in order to increase the abundance of the products above the detection threshold, we repeated the experiments by exposing the cultures to arachidonic acid (10μM), thus providing a substrate, during the 30-minute incubation period. In these cultures, 32 of the 48 compounds included in the panel were above the detection level (Table 1). Among the COX products, four exhibited significant increase following capsaicin treatment; PGD2, 8-iso-PGFa, 15dPGJ2 and 11d-tromboxane2 (11dTBX2; Figure 2H).

**Table 1.**
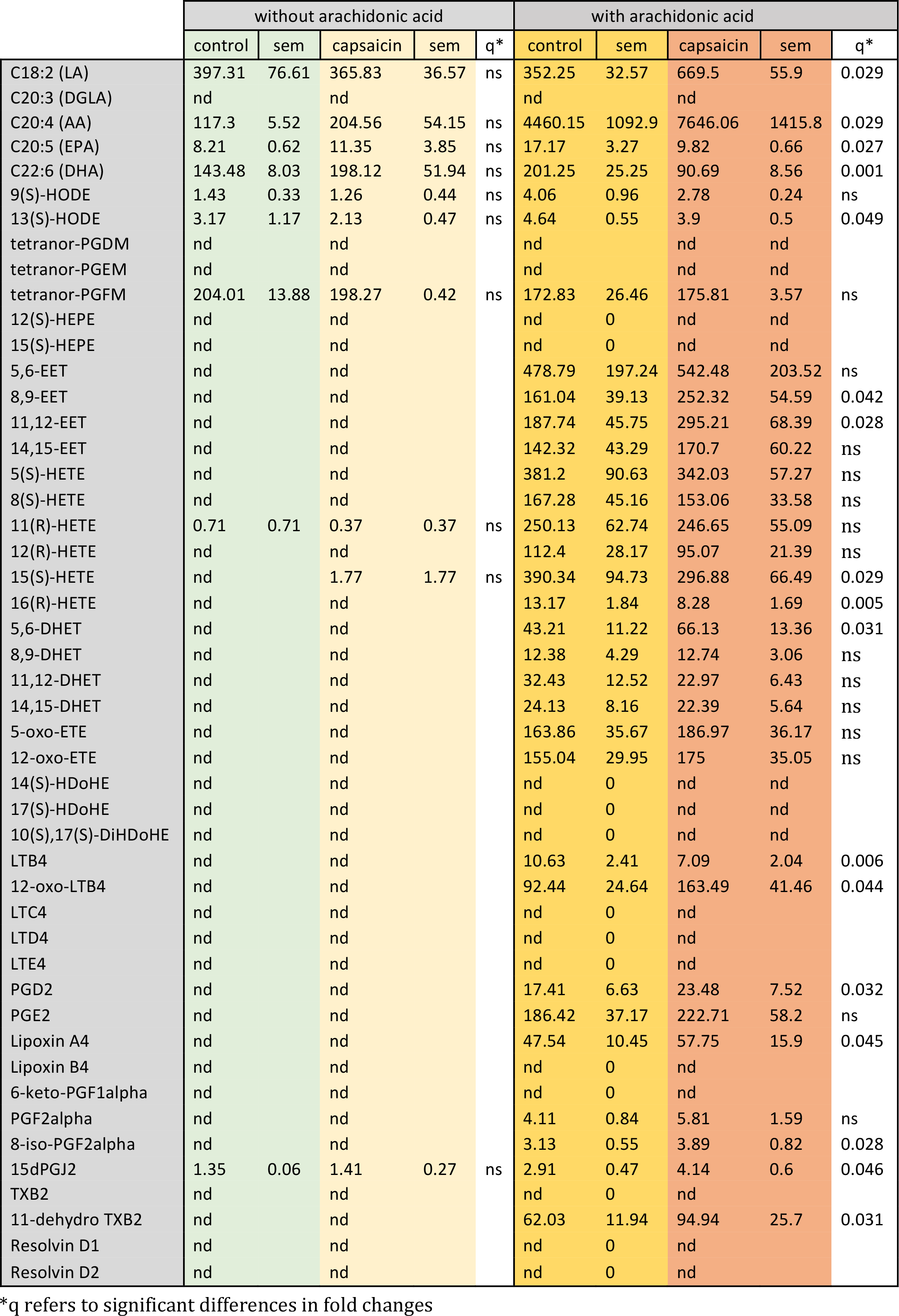
Concentration of 48 lipid mediators of inflammation in mouse primary sensory neuron cultures in naive condition and 30 minutes after incubating cells in capsaicin for 5 minutes (pg/ng protein).

## DISCUSSION

Eicosanoids and related compounds, including PGE2, 15dPGJ2 and PGD2, PGI2 have been implicated in inducing TRPV1 sensitisation (Moriyama *et al.*, 2005; Taiwo *et al.*, 1989; Taiwo & Levine, 1990). It is also reported that activation of primary sensory neurons leads to COX2 upregulation (Araldi *et al*, 2013; Fehrenbacher *et al*, 2005; Li *et al*, 2011; Samad *et al*, 2001). Further, it is reported that a group of primary sensory neurons exhibits sensitisation to capsaicin within 20-30 minutes after a previous capsaicin application (Zhang *et al.*, 2011). However, this is the first report which shows that increased *ptgs2* and COX2 expression and synthesis of a series of COX2 products are triggered by TRPV1 activation and thus underlie the development of sensitisation of TRPV1 in ~30 minutes after the first TRPV1 activation.

When capsaicin is repeatedly applied within a short period of time (i.e. tens of seconds – minutes), subsequent responses exhibit rapid desensitisation (for refs see Nagy *et al*., 2014). However, when capsaicin application is repeated only in tens of minutes, responses are sensitised (Zhang *et al.*, 2011). Here we report that, similarly to rat cultured primary sensory neurons (Zhang *et al.*, 2011), murine cultured primary sensory neurons also exhibit this type of sensitisation, though there are some differences between the two species. In mouse primary sensory neurons, the estimated proportion of cells exhibiting sensitisation are lower and the correlation between the amplitude of the first and second responses is less pronounced than in rat primary sensory neurons (Zhang *et al.*, 2011). Regarding the proportion of sensitizer neurons, however, it should be noted that our estimation maybe conservative as only neurons which exhibited at least a 5% increase in the amplitude of the second capsaicin-evoked responses were considered sensitizers. The assumption that we underestimated the proportion of sensitizers is supported by the findings that while ~1/3 of the capsaicin-responsive neurons exhibited at least 5% increase in responses to the second capsaicin application, about 1/2 of the TRPV1-expressing neurons appeared to express COX2 both in the control environment and after capsaicin exposure.

While Zhang and co-workers found involvement of ERK1/2 and CaMKII signalling (Zhang *et al.*, 2011), here we have identified further down-stream elements of the signalling pathway(s) which lead to capsaicin-evoked delayed sensitisation of TRPV1mediated responses. We found that blocking COX activity - either by the nonselective COX inhibitor indomethacin, or COX2 activity by the selective COX2 inhibitor NS-398 - almost completely eliminated the delayed sensitisation. These data suggest that COX2 activity is pivotal in delayed TRPV1 sensitisation. However, the few sensitizer neurons remaining in our samples after blocking COX or COX2 indicate that in addition to COX2, other mechanisms may also contribute to TRPV1 feed-forward sensitisation, albeit to a seemingly lesser extent than COX2.

Results of the RT-PCR, Western blotting and immunostaining together confirmed the role of COX2 in the capsaicin-induced delayed TRPV1 sensitisation as we detected upregulated synthesis of *ptgs2* and COX2. In accordance with findings that a group of primary sensory neurons expresses COX2 (Araldi *et al.*, 2013; Fehrenbacher *et al.*, 2005; Li *et al.*, 2011; Samad *et al.*, 2001), we found basal *ptgs2* and COX2 expression in our neuron samples. Our data indicate that COX2 basal expression, in agreement with previous reports on inducible COX2 expression in a group of primary sensory neurons (Fehrenbacher *et al.*, 2005; Li *et al.*, 2011), was significantly increased after capsaicin exposure. Further, here we report that capsaicin application to cultured primary sensory neurons also results in increased synthesis of COX2 products - though only when cells were presented by the COX2 substrate arachidonic acid. The failure in finding COX2 products with the absence of arachidonic acid in the buffer indicates the concentration of COX2 products without the presence of exogenous arachidonic acid is below the detection threshold when dissociated primary sensory neurons derived from one mouse are divided into control and treated cultures. Interestingly, in addition to several COX products, the concentration of several fatty acids and lipoxygenase products were also increased after arachidonic acid supplementation. Importantly, several of those upregulated lipoxygenase products, for example LTB4 or 15-(S)-HETE, might also contribute to delayed TRPV1 sensitisation; for example, in cells unaffected by the COX and COX2 inhibitors (Koskela *et al*, 2012; Zinn *et al*, 2017).

Several COX2 products including PGE2, 15dPGJ2 and PGD2, PGI2 have been implicated in acting on TRPV1-expressing primary sensory neurons (Moriyama *et al.*, 2005; Taiwo *et al.*, 1989; Taiwo & Levine, 1990). Our targeted UPLC-MS study revealed that 4 of the 7 COX products included in the panel we used (PGD2, PGF2a, 15dPGJ2 and 11dTBX2), showed significantly increased synthesis after capsaicin exposure of cultured murine primary sensory neurons. This finding suggests that enzymes downstream of COX2 may exhibit differential expression and/or activation in primary sensory neurons after capsaicin exposure.

Among the upregulated prostaglandins we found PGD2, which recently has been shown to activate PKA type II in TRPV1-expressing primary sensory neurons (Isensee *et al.*, 2014). As shown previously, PKA-mediated TRPV1 phosphorylation results in TRPV1 sensitisation (Rathee *et al.*, 2002) indicating that PGD2 might indeed sensitise TRPV1. However, Isensee and co-workers (Isensee *et al.*, 2014) found prostaglandin D synthase expression in primary sensory neurons that lack TRPV1 activity. Therefore, at present it is not clear whether, in our cultures, TRPV1expressing neurons themselves or non-TRPV1-expressing cells, activated by some agents released from TRPV1-expressing neurons by capsaicin, constitute the source of PGD2.

Another COX-related product we found upregulated, after exposing our cultures to capsaicin, is 15d-PGJ2. This product is synthesised from PGD2 through PGJ2 (Simmons *et al*, 2004). Recently, 15d-PGJ2 has been reported to activate TRPV1 through covalently binding to the ion channel (Shibata *et al*, 2016).

We also found PGI2 concentration upregulated by capsaicin exposure in our cultures. Importantly, PGI2 sensitises both heterologously expressed and native TRPV1 (Moriyama *et al.*, 2005). This PGI2-induced sensitisation contributes to the development of inflammatory heat hypersensitivity (Moriyama *et al.*, 2005).

Interestingly, even though the concentration of PGE2 (the best known, most powerful TRPV1 sensitizer, and widely studied COX product) was increased following capsaicin exposure - this change failed to reach the level of significance. This result might suggest that PGE2 has little role in the development of feed-forward TRPV1 sensitisation. Instead, PGE2 released from monocytes or differentiated macrophages (Sha *et al*, 2012) may have a greater contribution to a quick sensitisation of TRPV1 during inflammation (Grösch *et al*, 2017).

In summary, here we report the phenomenon of feed-forward TRPV1 sensitisation in murine primary sensory neurons, which depends largely on COX2 activity. We also report that COX2 activity is accompanied by upregulated synthesis of the enzyme itself and a significant increase in the synthesis of a number of COX-related products. While COX products released from immunocompetent cells sensitise TRPV1 seconds after acting on TRPV1-expressing primary sensory neurons, agents produced by COX2 in TRPV1-expressing primary sensory neurons themselves appear to prolong the sensitisation state. Hence, the COX2-mediated feed-forward TRPV1 sensitisation represents another ‘layer’ of PG-mediated TRPV1 sensitisation. Interestingly, our data suggest that COX2-dependent sensitisation through peripheral and primary sensory neuron mechanisms may involve different COX-related agents. The relative contribution of TRPV1 sensitisation through immunocompetent and primary sensory neuron mechanisms is not known at present. Further, the receptors involved in the delayed sensitisation process and the signalling pathways have not been elucidated. Nevertheless, these findings, in addition to the “immediate” (within minutes) and “late” (within hours) feed-forward TRPV1 sensitisation (mediated respectively by TRPV1 activators such as anandamide (Nagy *et al.*, 2009; van der Stelt *et al.*, 2005) and secondary gene products such as PARs (Chen *et al.*, 2013)) reveal a novel, delayed (~30-minute) mechanism of feed-forward autocrine signalling leading to TRPV1 sensitisation through the upregulation of immediate early gene and gene products. Therefore, these findings also indicate that, in addition to TRPV1 sensitisation by inflammatory mediators, feed-forward autocrine TRPV1 sensitisation (“autosensitisation”) is multifaceted and that COX2 inhibition in primary sensory neurons is important in reducing inflammatory heat hypersensitivity.

## MATERIALS AND METHODS

Experiments were conducted in accordance with guidelines published by the International Association for the Study of Pain (IASP)(Zimmermann, 1983), UK Animals (Scientific Procedures) Act 1986, revised National Institutes of Health *Guide for the Care and Use of Laboratory Animals*, Directive 2010/63/EU of the European Parliament and of the Council on the Protection of Animals Used for Scientific Purposes, Good Laboratory Practice and ARRIVE guidelines. A total of 24 wild type C57BL/6 mice were used. Mice were killed humanely following Isoflurane anaesthesia. Every effort was taken to minimize the number of animals.

### Culturing primary sensory neurons

Primary sensory neuron cultures were prepared as described (Nagy & Rang, 1999). Briefly, dorsal root ganglia (DRG) from the 3^rd^ cervical to the 6^th^ lumbar segments were harvested, placed into Ham’s Nutrient Mixture F12 supplemented with Ultraser G (2% final concentration), L-glutamine (0.2mM final concentration), penicillin (100IU/ml final concentration) and streptomycin (100μg/ml final concentration) and incubated at 37oC for 2.5 hours with collagenase type IV (312.5-625IU/ml final concentration) followed by an additional 30 minutes incubation in the presence of trypsin (2.5mg/ml final concentration). Ganglia were then washed in F12, triturated then plated onto poly-DL-ornithine-coated coverslips. Cultures were incubated in the supplemented F12 at 37oC for 24-48 hours before being used for further experiments.

### Calcium imaging

Imaging was conducted as described previously (Varga *et al*, 2014). Briefly, cells were loaded with 1μM FURA-2 (Invitrogen) and 2mM probenecid (Invitrogen) for 45-60 minutes in extracellular buffer (NaCl, 150mM; KCl, 5mM; CaCl_2_, 2mM; MgCl_2_, 2mM; glucose, 10mM and HEPES, 10mM; pH7.4) at 37oC. Coverslips were superfused (~1.5ml/minute) with the extracellular buffer at 37oC in a flow chamber. The outlet of the perfusion system was positioned next to the group of cells from which the recordings were obtained. Only one field of view was tested on each coverslip. Cells were exposed to excitation light alternating between the wavelengths of 355 and 380nM (OptoLED Light Source, Cairn Research, UK). The fluorescent signals emitted in response to each excitation were captured with an optiMOS camera (QImiging, Canada) connected to a PC running the WinFluor software package (John Dempster, Strathclyde University, UK).

Images were analysed offline with the WinFlour and Clampfit (Molecular devices, USA) software packages. Briefly, after establishing the regions of interest, the time course of changes in the fluorescent ratios were examined and exported to Clampfit where the standard deviation (SD) of the baseline activity and the maximum amplitudes of responses were measured. Change in the fluorescent ratio was accepted as a response if the generation of the response corresponded to the drug application and the amplitude of the change was greater than 3 times the SD of the baseline activity. The number of cells in various groups (i.e. sensitizer and nonsensitizer neurons) and the amplitude of responses of cells from at least three independent cultures were pooled and averaged.

### Reverse transcriptase polymerase chain reaction (RT-PCR)

RT-PCR was prepared as described previously (Varga *et al.*, 2014). Briefly, DRG cells were washed with Hank’s balanced salt solution (HBSS) buffer (pH 7.4; 138 mM NaCl; 5 mM KCl; 0.3 mM KH_2_PO_4_; 4 mM NaHCO_3_; 2 mM CaCl_2_; 1 mM MgCl2; 10 mM HEPES; 5.6 mM glucose) and incubated at 37.0°C in either 500 nM capsaicin solution (capsaicin-treated group) or HBSS (control group) for 5 minutes, followed by incubation in HBSS for 30 minutes. RNA was subsequently extracted using the RNeasy Plus Mini Kit (QIAGEN, UK) according to manufacturer’s instructions. RNA was reverse transcribed using Oligo(dT)12-18 primers (500 μg/ml; Invitrogen, UK), dNTP Mix (10 mM; Promega, UK), 5x First Strand Buffer (Invitrogen, UK) and 0.1 M DTT (Thermo Fisher Scientific, UK) and SuperScript II (Invitrogen, UK).

PCR for COX-2 and glyceraldehyde 3-phosphate dehydrogenase (GAPDH), as internal control, were performed using cDNA, 5x Green GoTaq® Reaction Buffer (Promega, UK), 25 mM magnesium chloride, PCR Nucleotide Mix (Promega, UK) and GoTaq® Flexi DNA Polymerase (Promega, UK). GAPDH primers were proprietary and supplied by Primerdesign Ltd, UK. Primers for COX-2 (NM_011198) were designed using PrimerBlast and are as follows: forward 5’TCCATTGACCAGAGCAGAGA −3’; reverse: 5’-TGGTCTCCCCAAAGATAGCA −3’. Optimal PCR annealing temperature for COX-2 was determined by gradient PCR annealing at a range from 53.0°C to 61.0°C. Each amplification of the 30 cycles consisted of 30 seconds at 95.0 °C, 1 minute at 58.0°C for COX2 or 60.0°C for GAPDH, and a final 1 minute at 72.0°C. Reactions were performed in thermal cycler Eppendorf Mastercycler Personal (Eppendorf, UK). PCR amplicons along with 100 bp DNA Ladder (New England Biolabs, UK) were visualised on 2% agarose gel using ethidium bromide. Images were captured under UV light using G:Box system (Syngene, UK) and processed with GeneSnap software (Syngene, UK). Digital image analysis of PCR amplicons was conducted using ImageJ software to quantify the intensity of PCR products, determined by the area encompassed by the intensity peaks. Level of COX2 transcription was normalised to the expression level of the housekeeping gene, GAPDH.

### Western blotting

Cells were incubated in 500nM capsaicin or in HBSS at 37°C at 37°C for 5 minutes then in HBSS supplemented with 5 μg/mL selective proteasome inhibitor MG-132 in DMSO supplemented for 30 mins. 50 μL of NP40 lysis buffer (Invitrogen, UK) supplemented with 1mM phenyl methane sulfonyl fluoride (PMSF) and 1× protease inhibitor cocktail (Sigma-Aldrich, USA) was used for protein extraction. The lysates were transferred to Eppendorf tubes and spun down at 14,800 rpm at 4°C for 10 minutes and the supernatant collected. The protein concentrations of cell lysates were measured before the SDS-PAGE. The supernatant was mixed with 6x reducing sample buffer (500 mM Tris pH6.8, 0.1 mg/mL SDS, 34% glycerol, 0.1mg/mL bromophenol blue, 5% β-mercaptoethanol) and heated at 98°C for 2 mins. An equal mass of proteins was loaded onto 9% polyacrylamide gels. Proteins were separated and transferred onto nitrocellulose membrane (GE Healthcare Life Sciences). The membrane was washed in tris-buffered saline containing 0.1% Tween (Sigma; (TBST) then blocked in 5% non-fat milk or bovine serum albumin (BSA; Sigma) in TBS-T at room temperature for 1 hour. The membrane was then probed with 1:2500 diluted anti-COX-2 (ab15191, Abcam) and 1:2000 diluted anti-β-tubulin III (neuronal) (T8578, Sigma-Aldrich), supplemented in 5 mL of 5% non-fat milk or BSA in TBS-T at 4°C overnight. The reaction was visualised using HRP-conjugated anti-rabbit IgG (7074S, Cell Signalling) and anti-mouse IgG (7076S, Cell Signalling) and the Pierce ECL 2 Western blotting substrate following the manufacturer’s instructions. The membranes were then imaged by the imaging machine GeneGnome XRQ SynGene. Analysis of immunoblots was performed using ImageJ. The intensity of the COX2 signal was normalised to the intensity of the β-tubulin III signal.

### Immunostaining

Cells were incubated in 500nM capsaicin or in HBSS at 37°C for 5 minutes, followed by an incubation in HBSS supplemented with 5 μg/mL selective proteasome inhibitor MG-132 in DMSO at 37°C for 30 minutes and fixing for 15 minutes in 4% paraformaldehyde. Cells were permeabilised (0.3% Titon X), blocked in 5% normal goat serum and incubated in anti-COX-2 M19 (sc-1747, Santa Cruz) antibody (1:200) for overnight incubation at 4°C. The reaction was visualised using an anti-Goat Cross-Adsorbed Alexa Fluor 488 (A-11055, Invitrogen; 1:1000) antibody and covered in Vectashield Hard Set Antifade Mounting Medium with DAPI (H-1500, Vector Laboratories). Images were taken using a Leica DMBL fluorescence microscope (Leica, Germany) with a Retiga 2000R digital camera (QImaging, USA) controlled by Leica Application Suite. All the images of randomly taken fields were taken with the same camera settings and conditions.

Immunofluorescence images were also analysed using ImageJ. The average COX2 and TRPV1 immunofluorescent signals were measured in a group of neurons in randomly selected areas in capsaicin-treated and control coverslips prepared from 3 cultures (>100 cells/culture). Intensity values from each coverslips were plotted, the threshold for positive immunofluorecent signals and the average signal intensity and the proportion of COX2- and/or TRPV1-expressing cells was established (Sousa-Valente *et al*, 2017).

### Measuring capsaicin-induced eicosanoid production

Cells were incubated at 37°C for 5 minutes in capsaicin (500nM) or HBSS buffer for control. The wells were then incubated for an additional 30 minutes in HBSS or in HBSS with 10μM arachidonic acid. Equivolume of 100% methanol was added to wells. Then, cells were scraped, collected in Eppendorf tubes, vortexed and 50μL of the lysate were removed from each sample for protein quantification. The samples were stored at −20°C.

Protein quantification was performed using the Pierce™ BCA protein assay kit (Thermo Fisher Scientific, UK) according to the instructions. Standards and samples were measured in duplicates. Reading was performed at a wavelength of 540nm in a BioTek™ ELx800™ spectrophotometer (Fisher Scientific, UK) and duplicate readings were averaged.

The rest of the samples were analysed at the MRC-NIHR National Phenome Centre (UK) using a novel targeted assay developed for the quantification of 48 lipid mediators of inflammation (Wolfer *et al*, 2015) . Briefly, separation was performed using a Waters HSS T3 column. Samples were then analysed in negative ion mode in a Waters Xevo TQ-S triple quadrupole mass spectrometer (MS) with an electrospray ionisation (ESI) source. Concentrations detected by UPLC-MS were first normalised to the protein content of the samples. Then, normalised compound concentrations of the capsaicin-treated cells were normalised to values established in the respective control.

### Statistical analysis

Normal distribution of data was assessed using the Shapiro-Wilk test. Significant change in the number of sensitising/non-sensitising and immunostained cells was assessed using Fischer’s exact test. Amplitudes in Ca^2+^ imaging were compared using Student’s t-test or ANOVA followed by Tukeys post-hoc test as appropriate. Changes in both expression values and the concentration of lipid mediators were compared using Student’s t-tests. In the case of the concentration of lipid mediators, Student’s ttest values were corrected for false discovery rate using the Benjamini-Hochberg approach. Values are expressed as average+/−standard error of mean (SEM). Differences were regarded as significant at p/q<0.05.

## ACKNOWLEDGEMENTS

JS-V and PLM have been supported, respectively by a PhD studentship from Fundacao para a Ciencia e a Tecnologia (Portugal) and a British Journal of Anaesthesia Research Grant.

The authors are grateful for Ms Berenike Buhl for her technical assistance.

## AUTHOR CONTRIBUTIONS

TL: calcium imaging

GW: Western-blotting, immunostaining, writing up

VCCH: RT-PCR, writing up

DS: UPLC-MS, writing up JVS: writing up

PLM: western-blotting, immunostaining

IN: conceptualisation, managing/leading project, writing up

## DATA AVAILABILITY

This study includes no data deposited in external repositories.

## EXTENDED VIEW FIGURE LEGENDS

**Extended View Figure 1.**
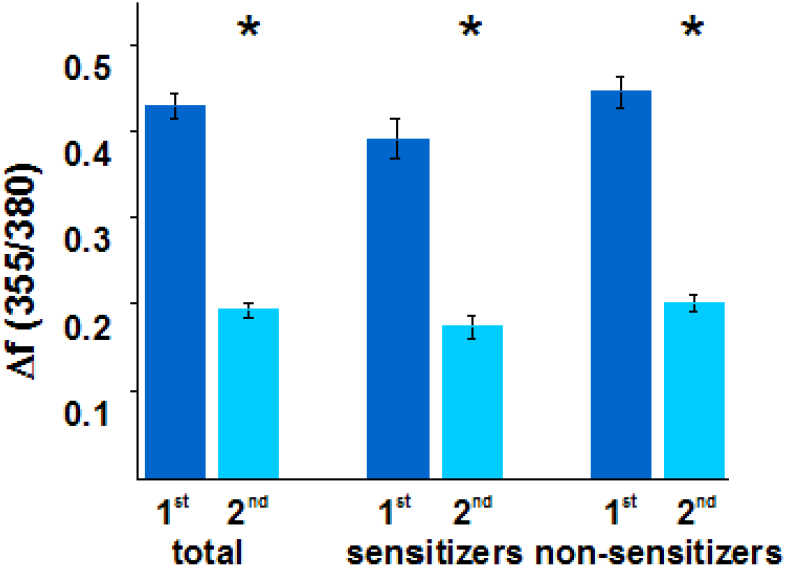
Amplitude of the first (1^st^) and second (2^nd^) KCl-evoked responses in all cultured murine primary sensory neurons (total), in sensitizer neurons (sensitizers) and non-sensitizer neurons (non-sensitizers) in control buffer. The second responses were significantly reduced in all groups (p<0.0001, Student’s t-test, n=288).

**Extended View Figure 2.**
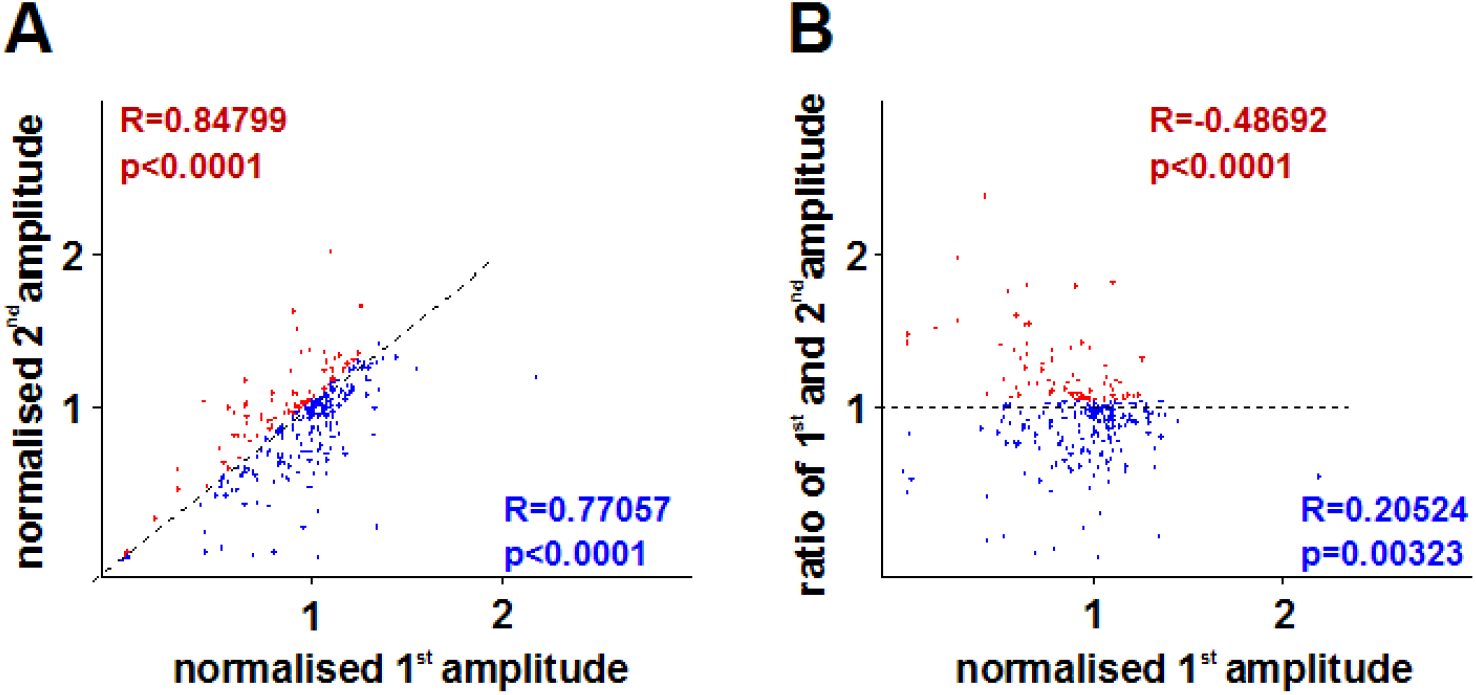
(A) Correlation between normalised capsaicin-evoked first and second responses of sensitizer (red) and non-sensitizer (blue) cells. In both group there is a significant correlation. Dotter line indicates first normalised response/second normalised response=1. (B) Correlation between the first normalised capsaicin-evoked response and the ratio of the first and second capsaicin-evoked responses in sensitizer (red) and non sensitizer (blue) cells. Sensitizers with smaller first responses tend to exhibit greater increase in the second response. Dotted line indicates a ratio of the first and second capsaicin-evoked responses=1.

**Extended view Figure 3.**
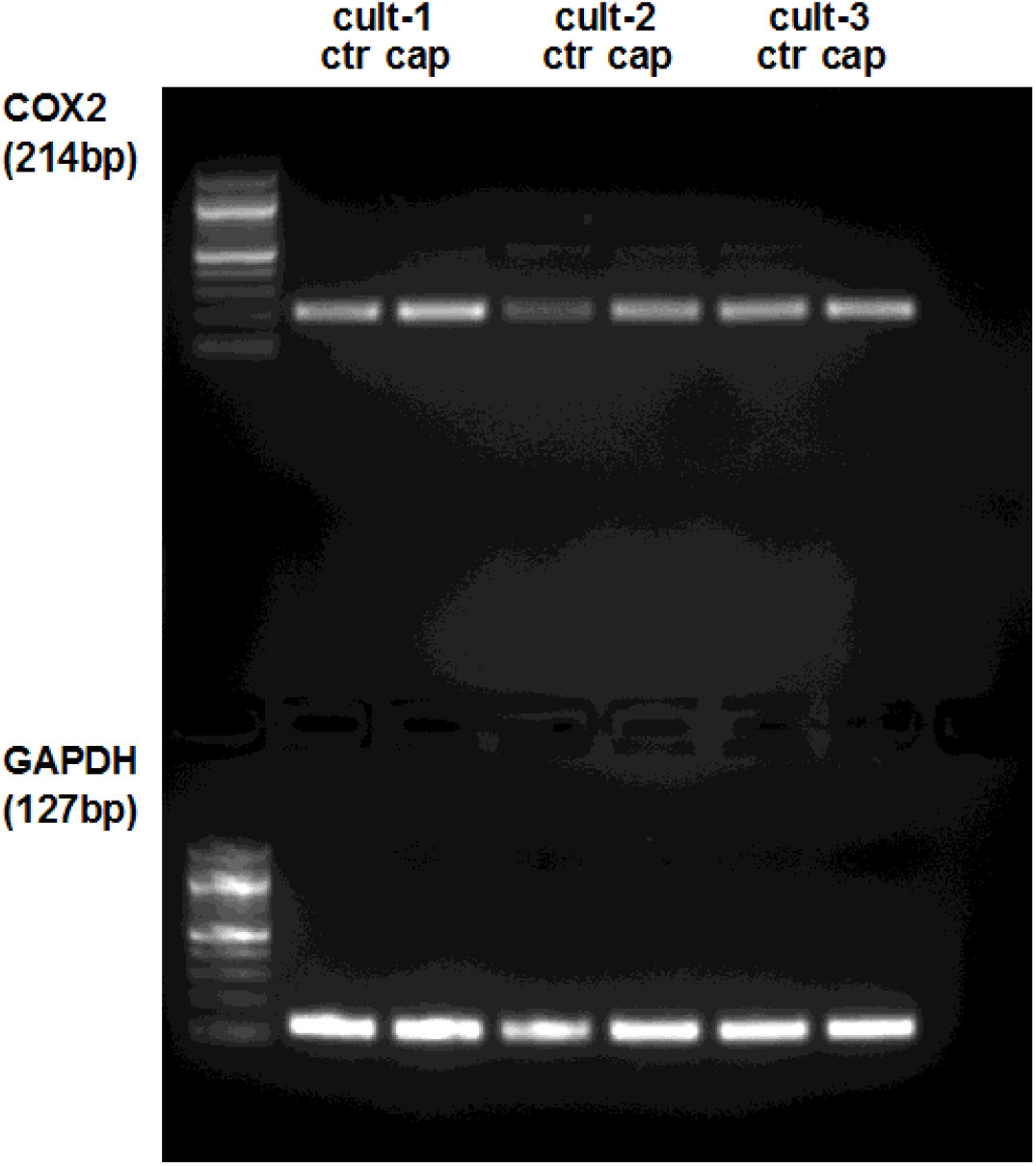
Full gel image of RT-PCR products amplified using primer pairs for murine *ptgs2* (COX2) and *gapdh* (GAPDH) and cDNA prepared using RNA isolated from three cultures (cult-1 – cult-3) of murine primary sensory neurons 25 minutes after incubating the cells in control (ctr) buffer or in the presence of 500nM capsaicin (cap) for 5 minutes.

**Extended View Figure 4.**
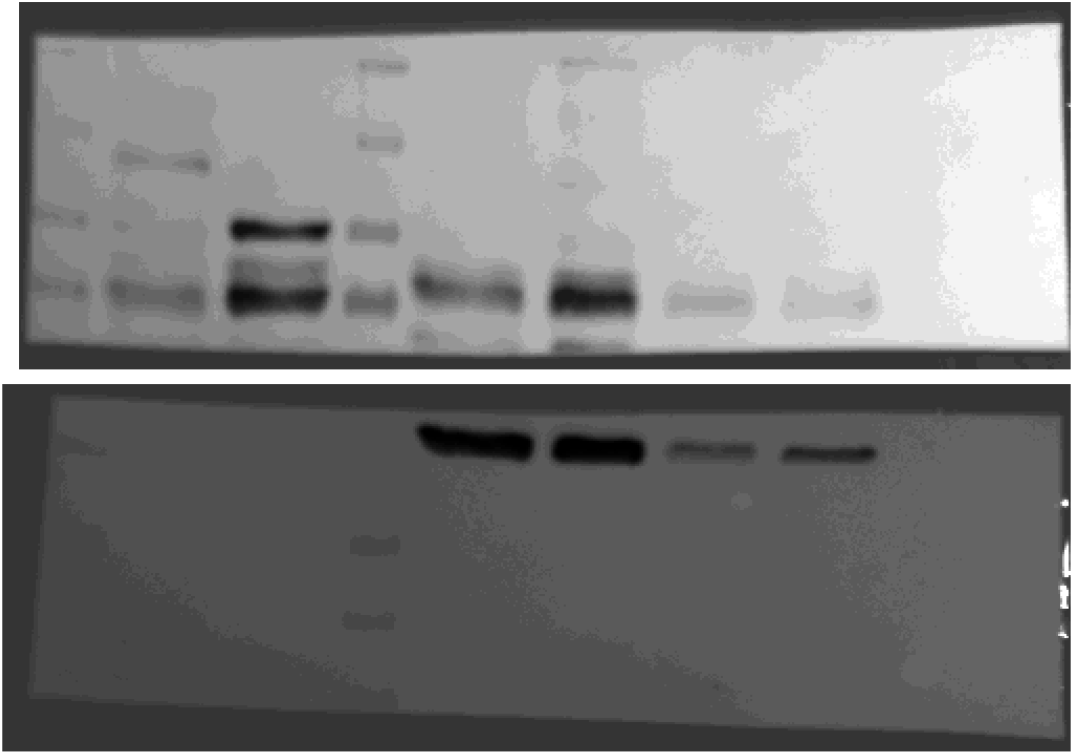
Full gel image of immunoblots prepared from protein samples extracted from culture 3. The culture was prepared from murine dorsal root ganglia. Proteins were extracted 25 minutes after incubating the cells in control buffer or in the presence of 500nM capsaicin for 5 minutes. Membranes were incubated in antibodies raised against COX2 (upper panel) and GAPDH (lower panel). Proteins were loaded at different concentrations. Lane 1: size marker, lane 2: control sample (highest concentration), lane 3: capsaicin-treated sample (highest concentration), lane 4: size marker, lane 5: control sample (intermediate concentration), lane 6: capsaicin-treated sample (intermediate concentration), lane 7: control sample (lowest concentration), lane 8: capsaicin-treated sample (lowest concentration). Blots of the intermediate concentrations were used for Figure preparation and quantification.

## REFERENCES

Amadesi S, Cottrell GS, Divino L, Chapman K, Grady EF, Bautista F, Karanjia R, Barajas-Lopez C, Vanner S, Vergnolle Net al (2006) Protease-activated receptor 2 sensitizes TRPV1 by protein kinase Cepsilon- and A-dependent mechanisms in rats and mice. J Physiol 575: 555–571

Araldi D, Ferrari LF, Lotufo CM, Vieira AS, Athié MC, Figueiredo JG, Duarte DB, Tambeli CH, Ferreira SH, Parada CA(2013) Peripheral inflammatory hyperalgesia depends on the COX increase in the dorsal root ganglion. Proceedings of the National Academy of Sciences of the United States of America 110: 3603–3608

Berta T, Qadri Y, Tan PH, Ji RR(2017) Targeting dorsal root ganglia and primary sensory neurons for the treatment of chronic pain. Expert Opin Ther Targets 21: 695–703

Caterina MJ, Leffler A, Malmberg AB, Martin WJ, Trafton J, Petersen-Zeitz KR, Koltzenburg M, Basbaum AI, Julius D(2000) Impaired nociception and pain sensation in mice lacking the capsaicin receptor. Science 288: 306–313

Caterina MJ, Schumacher MA, Tominaga M, Rosen TA, Levine JD, Julius D(1997) The capsaicin receptor: a heat-activated ion channel in the pain pathway. Nature 389: 816–824

Chen D, Wang Z, Zhang Z, Zhang R, Yu L(2013) Capsaicin up-regulates proteaseactivated receptor-4 mRNA and protein in primary cultured dorsal root ganglion neurons. Cellular and molecular neurobiology 33: 337–346

Fehrenbacher JC, Burkey TH, Nicol GD, Vasko MR(2005) Tumor necrosis factor alpha and interleukin-1beta stimulate the expression of cyclooxygenase II but do not alter prostaglandin E2 receptor mRNA levels in cultured dorsal root ganglia cells. Pain 113: 113–122

Fukuoka T, Tokunaga A, Tachibana T, Dai Y, Yamanaka H, Noguchi K(2002) VR1, but not P2X(3), increases in the spared L4 DRG in rats with L5 spinal nerve ligation. Pain 99: 111–120

Grösch S, Niederberger E, Geisslinger G(2017) Investigational drugs targeting the prostaglandin E2 signaling pathway for the treatment of inflammatory pain. Expert opinion on investigational drugs 26: 51–61

Isensee J, Wenzel C, Buschow R, Weissmann R, Kuss AW, Hucho T(2014) Subgroup-elimination transcriptomics identifies signaling proteins that define subclasses of TRPV1-positive neurons and a novel paracrine circuit. PLoS One 9: e115731

Ji RR, Samad TA, Jin SX, Schmoll R, Woolf CJ(2002) p38 MAPK activation by NGF in primary sensory neurons after inflammation increases TRPV1 levels and maintains heat hyperalgesia. Neuron 36: 57–68

Kao DJ, Li AH, Chen JC, Luo RS, Chen YL, Lu JC, Wang HL(2012) CC chemokine ligand 2 upregulates the current density and expression of TRPV1 channels and Nav1.8 sodium channels in dorsal root ganglion neurons. Journal of neuroinflammation 9: 189

Khan AA, Diogenes A, Jeske NA, Henry MA, Akopian A, Hargreaves KM(2008) Tumor necrosis factor alpha enhances the sensitivity of rat trigeminal neurons to capsaicin. Neuroscience 155: 503–509

Koskela H, Purokivi M, Nieminen R, Moilanen E(2012) The cough receptor TRPV1 agonists 15(S)-HETE and LTB4 in the cough response to hypertonicity. Inflammation & allergy drug targets 11: 102–108

Li X, Kim JS, van Wijnen AJ, Im HJ(2011) Osteoarthritic tissues modulate functional properties of sensory neurons associated with symptomatic OA pain. Mol Biol Rep 38: 5335–5339

Moriyama T, Higashi T, Togashi K, Iida T, Segi E, Sugimoto Y, Tominaga T, Narumiya S, Tominaga M(2005) Sensitization of TRPV1 by EP1 and IP reveals peripheral nociceptive mechanism of prostaglandins. Molecular pain 1: 3

Nagy I (2004) Sensory processing: primary afferent neurons/DRG. Anesthetic Pharmacology: Physiologic Principles and Clinical Practice, eds: Evers and Maze: 187–197

Nagy I, Fedonidis C, Paule CC, Wahba J, Andrew P, Austin J, Yaqoob M, Buluwela L, Nagy B, Nyilas R(2009) NAPE-PLD is involved in anandamide synthesis in capsaicin-sensitive primary sensory neurons. J Physiol Sci 59: 422–422

Nagy I, Friston D, Torres J, Sousa-Valente J, Andreou A(2014) Pharmacology of the capsaicin receptor, transient receptor potential vanilloid type-1 ion channel. In: Progress in Drug Research: Capsaicin as a therapeutic molecule, eds: Abdel-Salam, Springer, Basel, pp. 39–76.

Nagy I, Rang HP(1999) Similarities and differences between the responses of rat sensory neurons to noxious heat and capsaicin. The Journal of neuroscience: the official journal of the Society for Neuroscience 19: 10647–10655

Placek K, Schultze JL, Aschenbrenner AC(2019) Epigenetic reprogramming of immune cells in injury, repair, and resolution. J Clin Invest 129: 2994–3005

Premkumar LS, Ahern GP(2000) Induction of vanilloid receptor channel activity by protein kinase C. Nature 408: 985–990

Rathee PK, Distler C, Obreja O, Neuhuber W, Wang GK, Wang SY, Nau C, Kress M(2002) PKA/AKAP/VR-1 module: A common link of Gs-mediated signaling to thermal hyperalgesia. J Neurosci 22: 4740–4745

Samad TA, Moore KA, Sapirstein A, Billet S, Allchorne A, Poole S, Bonventre JV, Woolf CJ(2001) Interleukin-1beta-mediated induction of Cox-2 in the CNS contributes to inflammatory pain hypersensitivity. Nature 410: 471–475

Sha W, Brüne B, Weigert A(2012) The multi-faceted roles of prostaglandin E2 in cancer-infiltrating mononuclear phagocyte biology. Immunobiology 217: 1225–1232

Shibata T, Takahashi K, Matsubara Y, Inuzuka E, Nakashima F, Takahashi N, Kozai D, Mori Y, Uchida K(2016) Identification of a prostaglandin D2 metabolite as a neuritogenesis enhancer targeting the TRPV1 ion channel. Sci Rep 6: 21261

Simmons DL, Botting RM, Hla T(2004) Cyclooxygenase isozymes: the biology of prostaglandin synthesis and inhibition. Pharmacological reviews 56: 387–437

Sousa-Valente J, Varga A, Torres-Perez JV, Jenes A, Wahba J, Mackie K, Cravatt B, Ueda N, Tsuboi K, Santha Pet al (2017) Inflammation of peripheral tissues and injury to peripheral nerves induce differing effects in the expression of the calcium-sensitive N-arachydonoylethanolamine-synthesizing enzyme and related molecules in rat primary sensory neurons. J Comp Neurol 525: 1778–1796

Taiwo YO, Bjerknes LK, Goetzl EJ, Levine JD(1989) Mediation of primary afferent peripheral hyperalgesia by the cAMP second messenger system. Neuroscience 32: 577–580

Taiwo YO, Levine JD(1990) Effects of cyclooxygenase products of arachidonic acid metabolism on cutaneous nociceptive threshold in the rat. Brain Res 537: 372–374

Van Buren JJ, Bhat S, Rotello R, Pauza ME, Premkumar LS(2005) Sensitization and translocation of TRPV1 by insulin and IGF-I. Molecular pain 1: 17

van der Stelt M, Trevisani M, Vellani V, De Petrocellis L, Schiano Moriello A, Campi B, McNaughton P, Geppetti P, Di Marzo V (2005) Anandamide acts as an intracellular messenger amplifying Ca^2+^ influx via TRPV1 channels. EMBO J 24: 3026–3037

Varga A, Jenes A, Marczylo TH, Sousa-Valente J, Chen J, Austin J, Selvarajah S, Piscitelli F, Andreou AP, Taylor AHet al (2014) Anandamide produced by Ca(2+)-insensitive enzymes induces excitation in primary sensory neurons. Pflugers Arch 466: 1421–1435

Wolfer AM, Gaudin M, Taylor-Robinson SD, Holmes E, Nicholson JK(2015) Development and Validation of a High-Throughput Ultrahigh-Performance Liquid Chromatography-Mass Spectrometry Approach for Screening of Oxylipins and Their Precursors. Anal Chem 87: 11721–11731

Woolf CJ, Ma Q(2007) Nociceptors--noxious stimulus detectors. Neuron 55: 353364

Yekkirala AS, Roberson DP, Bean BP, Woolf CJ(2017) Breaking barriers to novel analgesic drug development. Nature reviews Drug discovery 16: 545–564

Zhang X, Daugherty SL, de Groat WC (2011) Activation of CaMKII and ERK1/2 contributes to the time-dependent potentiation of Ca^2+^ response elicited by repeated application of capsaicin in rat DRG neurons. Am J Physiol Regul Integr Comp Physiol 300: R644–654

Zhang X, Huang J, McNaughton PA(2005) NGF rapidly increases membrane expression of TRPV1 heat-gated ion channels. EMBO J 24: 4211–4223

Zimmermann M (1983) Ethical guidelines for investigations of experimental pain in conscious animals. Pain 16: 109–110

Zinn S, Sisignano M, Kern K, Pierre S, Tunaru S, Jordan H, Suo J, Treutlein EM, Angioni C, Ferreiros Net al (2017) The leukotriene B4 receptors BLT1 and BLT2 form an antagonistic sensitizing system in peripheral sensory neurons. The Journal of biological chemistry 292: 6123–6134

